# Balanced Functional Module Detection in Genomic Data

**DOI:** 10.1101/2020.11.30.404038

**Authors:** David Tritchler, Lorin M Towle-Miller, Jeffrey C Miecznikowski

## Abstract

High dimensional genomic data can be analyzed to understand the effects of multiple variables on a target variable such as a clinical outcome, risk factor or diagnosis. Of special interest are functional modules, cooperating sets of variables affecting the target. Graphical models of various types are often useful for characterizing such networks of variables. In other applications such as social networks, the concept of balance in undirected signed graphs characterizes the consistency of associations within the network. To extend this concept to applications where a set of predictor variables influences an outcome variable, we define balance for functional modules. This property specifies that the module variables have a joint effect on the target outcome with no internal conflict, an efficiency that evolution may use for selection in biological networks. We show that for this class of graphs, observed correlations directly reflect paths in the underlying graph. Consequences of the balance property are exploited to implement a new module discovery algorithm, bFMD, which selects a subset of variables from highdimensional data that compose a balanced functional module. Our bFMD algorithm performed favorably in simulations as compared to other module detection methods that do not consider balance properties. Additionally, bFMD detected interpretable results in a real application for RNA-seq data obtained from The Cancer Genome Atlas (TCGA) for Uterine Corpus Endometrial Carcinoma using the percentage of tumor invasion as the target outcome of interest. bFMD detects sparse sets of variables within highdimensional datasets such that interpretability may be favorable as compared to other similar methods by leveraging balance properties used in other graphical applications.

## 1 Introduction

Clinical outcomes of interest, such as risk factors, disease diagnosis, or treatment success may be better understood by identifying associated genes within high-dimensional genomic datasets. Additionally, these associated genes should be associated with one another, such that they may form a “module” or “pathway” that coordinate together to ultimately associate with the targeted outcome of interest. We denote these outcome associated sets of variables as “functional modules”. Many methods currently exist to identify functional modules, but these existing functional module detection methods fail to consider the concept of “balance”, oftentimes used in other applications such as social networks. Balance assures no contradictions exist between variables within the functional module. For example, suppose the expression of a certain module gene promotes an increase in the outcome variable while another gene in the same module supresses it; balance says that the two genes must be negatively correlated; otherwise they would conflict with respect to their effect on the outcome.

Since observed correlations in a balanced sign graph directly reflect all paths in the underlying graph, we can use the observed correlation matrix to determine if a subset of the high dimensional set of variables satisfies the balance property. This leads to our new module discovery algorithm, bFMD. After the module variables are selected from the high dimensional data set, various analytical methods may be used for obtaining the graphical structure in detail. For example, modelling predictors as an undirected graph whose vertices influence the target is related to traceable regressions (Wermuth (2012)). The traceable regression analysis of our semi-directed graph is partitioned into an undirected graphical model analysis (eg. Edwards (2000), Edwards et al. (2010)) of the predictors and individual regressions of the target on each predictor (Wermuth and Cox (2013)). Such analyses are often difficult or impossible for high dimensional data, so dimension reduction prior to a fully detailed traceable regression analysis may be necessary (Bourgon et al. (2010), Towle-Miller et al. (2020)).

The concepts presented in this manuscript build on the concepts utilized in (Miecznikowski et al. (2016)) with added theoretical details and extensions using sparse principal components analysis for variable selection. Section 2 of this paper provides results on weighted undirected graphs which will be used in later sections of this manuscript. Importantly, since the weights of graph edges can be positive or negative, these are considered to be signed graphs (Chartrand (1977), ch.8). Section 3 builds on Section 2 by adding an outcome variable *Y* and associated directed edges to the graphical model, resulting in a semi-directed graph which we refer to as a functional module. In Section 4 we introduce the concept of balance in an undirected graph as described in Definition 1, and Definition 2 extends the concept to the semi-directed graphs which is of primary interest. Theorem 4.1 in Section 4 establishes a useful property using triplets of paths in a functional module. In section 5 we model the undirected subgraph as a Gaussian graphical model, we state Theorem 5.2 which relates observed covariances to the underlying graph, and then we state Theorem 5.3 which extends these results to the semi-directed case. Section 6 defines a matrix formed from covariances where the submatrix corresponding to the functional module must be positive in order to satisfy balance, as proven by Theorem 6.1. Section 7 introduces Balanced Functional Moledule Detection (bFMD), a new method for module discovery based on Theorem 6.1 and Perron theory of positive matrices. Section 8 presents Theorem 8.2 which motivates a metric that can be used to gage the balance of a computed module. Section 9 describes the simulation to be used and compares our method to existing methods. Section 10 applies bFMD to real world RNA-seq data obtained from The Cancer Genome Atlas (TCGA) (Tomczak et al. (2015)). The final section is a summary and discusses further points.

## 2 Preliminaries

Let *G* = (*X,E,w*) denote a weighted, undirected graph with vertex set *X* = {*x*_1_,*x*_2_,…,*x_p_*}; edge set E and weight function w: E → R. Note that *G* is a signed graph since the weights can be either negative or nonnegative. To alleviate cumbersome notation, we will interchangeably denote *x_i_* as *i* when considering *x_i_* as a graph vertex. We further denote the weight of an *i – j* edge as *w_ij_*.

We assume that a path *γ*: *i → j* of length *m* between vertices *i* and *j* creates an ordered sequence of non-repeated adjacent indices of the form *γ* = ((*i, k*_1_), (*k*_1_, *k*_2_),…, (*k_m_,j*)) ∈ *E^m^*, where *k_l_* corresponds to the intermitant indices between *i* and *j*. We calculate the path weight as *w^γ^* = *w*_*ik*1_ *w*_*k*_1_*k*_2__… *w_k_m_j_*. Note that the resulting sign of *w^γ^* will be negative if and only if there is an odd number of negative edges along the path. We additionally define a “walk” as a path with repeated vertices, and a “cycle” as a path where *i* = *j*.

Further note that we may consider the vertex set *X* as a vector or *x_i_* as a graph vertex/variable where appropriate when discussing the statistical properties throughout this manuscript.

## 3 Functional modules

We informally define a “functional module” to be a set of coordinated variables *X* = {*x_i_*; *i* = 1, 2,…*p*} which influence a outcome variable *Y*. Each variable in *X* influences *Y* either directly or indirectly by operating through other variables in the set *X*. Such modules that help describe the response *Y* can be profitably described by an underlying weighted undirected graph *G* = (*X, E, w*) under special properties.

*G* and *Y* create a semidirected graph *G ∪ Y*, and the vertices in G are each a potential predictor of *Y*. The directed portion of *G ∪ Y* corresponds to the edges connecting from *G* to *Y*, and the undirected portion of *G ∪ Y* corresponds to the edges that connect variables within *X* but not *Y*. We represent a semidirected path *α*: *i → Y* in *G ∪ Y* as two segments: an undirected path *γ*: *i – j* from *i* ∈ *G* terminating in some *j* ∈ *G* followed by a directed edge from *j* to *Y*. We define the weight of the path *α* from *i* to *Y* along *γ* to be 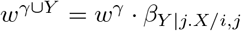, where *β_Y|j.X/i,j_* represents the partial regression coefficient of *j* in the regression model for response *Y* including all variables in *X except x_i_* and *w^γ^* represents the path weight from *i* to *j* in *G*. Note that *sign*(*α*) = *sign*(*w^γ^*) * *sign*(*β_Y|j.X/i,j_*) since *β_Y|j.X/i,j_* characterizes the directed edge, and in the special case when a path consists of only a directed edge from *x_j_* to *Y* the path weight equals *β_Y|j,X/j_*.

## 4 Balance in functional modules

Building on the definitions described in previous sections, we further require the signed graph G to be *balanced* (Harary (n.d.)). A characterization of balance which best results in desired properties of a functional module is described in Definition 4.1. For example, if i is related to j by a path of module elements, no other *i — j* paths in the network should exist with opposing signs. This balanced mechanism ensures no conflicts, and it greatly improves the interpretability in biological systems by identifying sets of variables which operate in this cooperative fashion.

### Definition 4.1 Balanced Sign Graph

*A signed graph G is “balanced” if and only if for every two vertices of G, all paths joining them have the same sign (Chartrand (1977), Ch. 8; Harary (n.d.)).*

To model our functional network, we extend the concept of balance to semi-directed graphs by requiring all semi-directed paths *α: i → Y* to have the same sign, which results in a similar interpretation to balance within *G*. More precisely, we require a functional network *G ∪ Y* to have the properties described in Definition 4.2. In summary, *G* is a *module* or network (by the second condition) with no internal conflict (by the first and fourth conditions), and the module affects *Y* (by the third condition).

### Definition 4.2 Balanced Functional Network (*G ∪ Y*)

*When the following conditions are satisfied, we say that G ∪ Y is a “balanced functional module”.*

1. *G is balanced.*
2. *G is connected: at least one path connecting each pair of elements within G exists.*
3. *Every vertex of G is connected to Y by a semi-directed path with at least one directed edge.*
4. *For a given vertex i ∈ G, all i — Y paths have the same sign.*

In Theorem 4.1, we establish a consequent constraint on triplets of paths in a balanced functional network.

### Theorem 4.1

*For balanced functional network G ∪ Y, let α ∪ Y equal a semi-directed i → Y path, γ equal an i — j path in G, and τ ∪ Y equal a semi-directed j → Y path. Then w^α∪Y^ · w^γ^ · w^τU∪Y^* > 0.

**Proof 4.1** *Note that w^γ^ · β_Y|j.X/j_ equals the weight of an i — Y path, so sign(w^α∪Y^*) = *sign(w^γ^ · β_Y|j.X/j_*) = *sign*(*w^γ^*) · *sign*(*β_y|j.X/j_*). *However, β_Y|j.X/j_ equals the weight of the directed j → Y edge which is also a j — Y path. Hence, sign(β_y|j.X/j_*) = *sign(w^τ∪Y^*). *Now, sign(w^α∪Y^*) = *sign(w^γ^) · sign*(*w^τ∪Y^*), *and the product of the three terms w^α∪Y^,w^γ^,w^τ∪Y^ must be positive.*

## 5 Statistical Model

We represent *G* statistically as a Gaussian undirected graphical model with a nonsingular concentration matrix Θ = Σ^-1^. The properties of Θ will mimic those from previous studies (Wermuth (1980), Whittaker (2009), Anufriev and Panvhenko (2015), Sulaimanov and Koeppl (2016)). Specifically, we denote the diagonal of Θ as *D* = *diag*(Θ_11_, Θ_22_,…, Θ_pp_), where *d_i_* = Θ_*ii*_ = *Var*(*x_i_*\*X*/*i*)^-1^ denotes the reciprocal of the partial variance of *x_i_*. Additionally, letting *I* denote an identity matrix of appropriate size, the adjacency matrix *A* = *I* — *D*^-1/2^Θ*D*^-1/2^ contains partial correlations with zeros on its diagonal, making 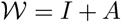 a matrix of partial correlations. Note that the off-diagonal elements of *A* correspond to the edge weights from graph *G*. From this we see that the partial covariance matrix is 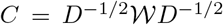, and the matrix of least square partial regression coefficients is 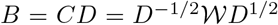. The covariance matrix Θ^-1^ = Σ = (*σ_ij_*) relates to the adjacency matrix *A* of the graph *G* through the expresion Σ = *D*^-1/2^(*I — A*)^-1^*D*^-1/2^ = *D*^-1/2^(*D*^-1/2^Θ*D*_-1/2_)^-1^*D*^-1/2^ = *D*^-1/2^(*D*^1/2^Σ*D*^1/2^)*D*^-1/2^.

Fully describing the undirected graph *G* requires Σ^-1^, which involves a large number of parameters and often does not exist for high dimensional data. Also, although the variables involved in the module are typically unknown in real applications, more detailed analyses may be subsequently performed after identifying the subset of module variables. Additionally, prior biological knowledge of the individual module elements may be sufficient to gain scientific understanding.

The covariance structure of *G* can be described in terms of a closure matrix *G*\* (Sulaimanov and Koeppl (2016), Meyer (2000) ch. 7). Powers of the adjacency matrix sum walks in *G* of a specific length, e.g. 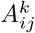 equals the sum of all *i — j* walk weights in *G* of length *k*. Then the Neumann series 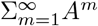 quantifies the connectivity of *G* and converges to (*I — A*)^-1^ = *D*^-1/2^Σ*D*^-1/2^ (Sulaimanov and Koeppl (2016)). The off-diagonal elements of Σ are then given by 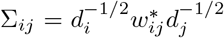, where each 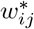 corresponds to a weight in the closure *G*\* of *A*, making Σ_*ij*_ the scaled sum of weights of all *i — j* walks in *G*. The Neumann series converges when the spectral radius of *A*, denoted *σ*(*A*), is less than 1, and 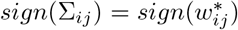 when the Neumann series converges. It may be shown then when all row sums of the absolute values in A are less than 1, then *σ*(*A*) < 1. This is interpreted as moderate conditional dependence among the variables since A consists of partial correlations (Anufriev and Panvhenko (2015)).

Balance is defined in terms of paths in the literature, while the preceding development describes Σ using walks. To bridge this disconnect we require Lemma 5.1.

### Lemma 5.1

*For a balanced graph G, the sign of an i — j walk is the sign of every i — j path in G.*

**Proof 5.1** *Let* γ *be an i — j walk in G. If γ is not a path then there exists a cycle where a repeated vertex is connected by a path within γ. Since every cycle in a balanced graph is positive (Harary (n.d.); Chartrand (1977), Theorem 8.2), the cycle can be excised from the walk without changing the sign of the walk. This process can be continued until ultimately an i — j path with the same sign as γ remains.*

Note that Lemma 5.1 implies that every *i — j* walk contains the same sign because according to Definition 4.1, every *i — j* path must have the same sign to be considered balanced. An immediate consequence is shown in Theorem 5.2.

### Theorem 5.2

*For a balanced graph G and corresponding covariance matrix* Σ = (*σ_ij_*), *if we assume moderate conditional dependence then the sign of σ_ij_ is equivalent to the sign of every i — j path in G.*

**Proof 5.2** *The sign of σ_ij_ equals the sign of the summed i — j walk weights in G, and each i — j path contains the same sign as the i — j walks.*

We extend the above concept to the semidirected case in Theorem 5.3.

### Theorem 5.3

*For a balanced functional network G ∪ Y with G satisying the conditions of Theorem 5.2:*

1. *Cov(i, Y) is a positively weighted sum of all i — Y walk weights*
2. *The sign of Cov(i, Y) equals the sign of every i — Y path.*
3. *Cov(i,Y) ≠ 0.*

**Proof 5.3** *By Proposition 5.5.1 of (Whittaker (2009)):*

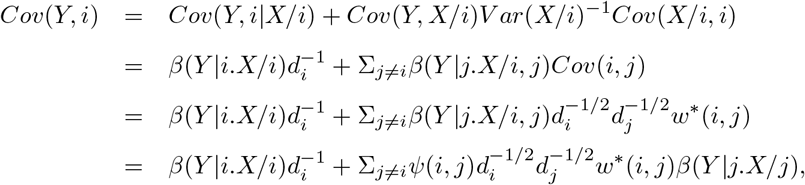

*where ψ*(*i, j*) = *β*(*Y|j.X/i, j*)/*β*(*Y|j.X/j*) *corresponds to the ratio of the partial regression coefficient for x- in the regression model with predictor set X/i to the coefficient for x- in the model including x_i_ as a predictor. The dk weights are positive since they are variances. ψ(i, j) is positive if adding *x_i_* to the predictors does not change the sign of the regression coefficient for x_i_. We take the condition of moderate conditional dependence required for Theorem 5.2 to imply that such sign change of coefficients will not occur. The first term of the last equation is proportional to the direct i → Y edge, and the summation in that equation is a positively weighted sum of indirect i → Y walks passing through other vertices in G. This proves the first statement of the theorem. By Lemma 5.1, w*(i, j) equals the sum of walk weights with the same sign as every i — j path, so the sign of w*\*(*i,j*)*β*(*Y|j.X/j*) *equals the sign of any i — Y path with weight w*(*i,j*)*β*(*Y|j.X/j*). *This proves the second statement. The third statement immediately follows because Cov*(*i, Y*) = 0 *implies that the sign of every i — Y path is zero, contradicting statement 3 of Definition f.2.*

## 6 Identification of module elements

We use Theorems 4.1, 5.2, and 5.3 to guide the discovery of subsets within X which influence Y and behave as a balanced graph. To this end, we define the matrix M in Theorem 6.1, and balanced functional modules will appear as positive submatrices of this matrix.

### Theorem 6.1

*If the graph G ∪ Y within X is a balanced functional module, then*

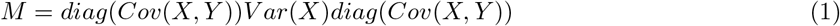

*is a positive matrix (i.e. all elements of M are > 0*).

**Proof 6.1** *By Theorems 5.2 and 5.3, sign(Cov(x_i_, Y)) equals the sign of any i — Y path α, sign(Cov(x-,Y)) equals the sign of any j — Y path τ, and sign(Cov(i, j)) equals the sign of any i — j path γ in G. Thus, for every i, j pair*, 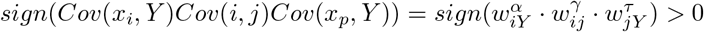 *by Theorem j.1.*

Note that the (*i,j*)th element of *M* equals *Cov*(*x_i_,Y*)*Cov*(*i, j*)*Cov*(*x_p_, Y*), making it large when *i* and *j* are highly correlated with each other and with *Y*. Theorem 6.1 provides a mechanism for identifying the subset of module elements by analyzing a matrix similar to (1) by using the full data collected. The data collected will contain a high dimensional set of variables, and the module *M* will be a subset of the larger matrix. If we compute the matrix *M*\* = *diag*(*Cov*(*X*, Y*))Σ(*X*\*)*diag*(*Cov*(*X*,Y*)) for entire set of predictors *X*\*, then the balanced functional module *M* will appear as a positive submatrix of *M*\*. The nonzero elements correspond to correlated variable pairs that are additionally associated with *Y*. When elements equal zero, they fail to satisfy all three conditions. Since the ordering of the variables is arbitrary, we can write *M*\* as a matrix composed of zeros with a positive upper diagonal block *M* corresponding to the functional module. We then model the observed data matrix as *W* = *M*\* + *E*, where *E* is a matrix of mean zero errors. Thus, the identification of a positive submatrix within *W* can approximate the functional module *M*.

To depict this, note that *W* can be written as the Hadamard product

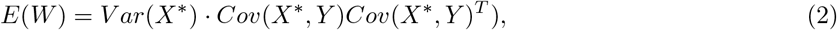

where the operator o denotes element-wise multiplication. The left matrix, *Var*(*X*\*), contains the information on the modular structure, and the right matrix, *Cov*(*X*,Y*)^*T*^), is positive when the variables corresponding to that element are correlated with eachother and with *Y*. Taken from Miecznikowski et al. (2016), Figure 1 visually depicts the formation of the Hadamard product (2) for 3 simulated modules, one of which is functional.

**Figure 1:**
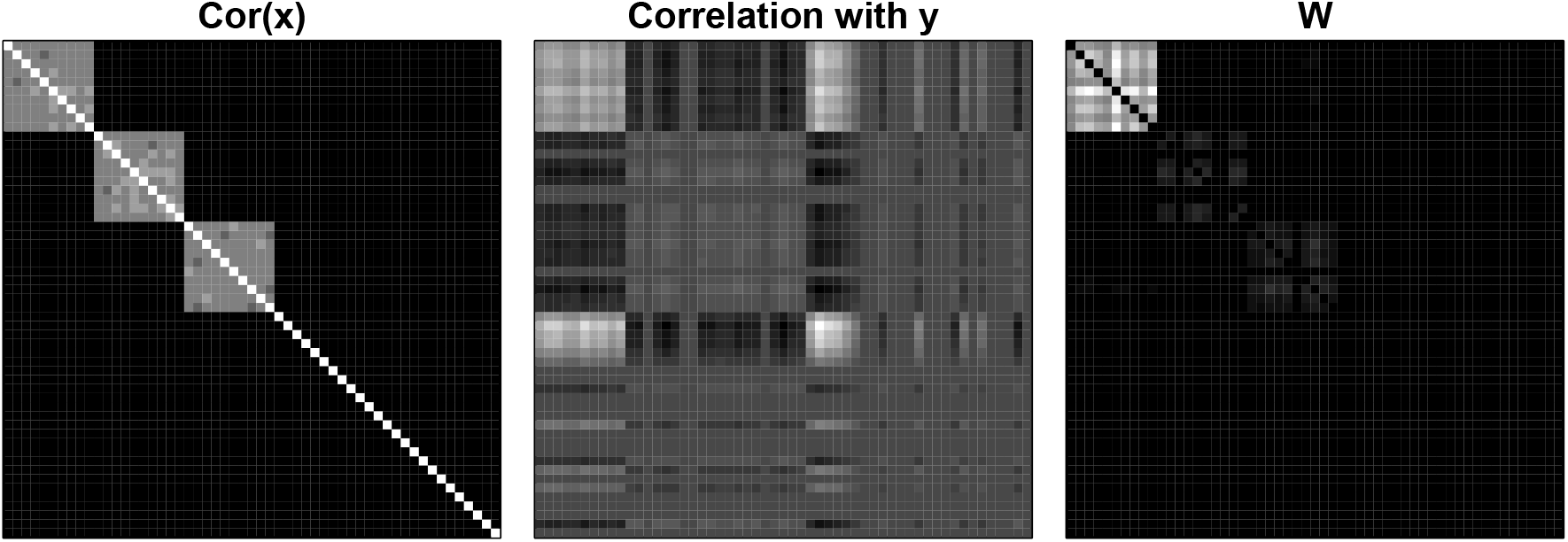
The formation of *W* = *Cov*(*X*) · (*Cov*(*X, Y*)*Cov*(*X,Y*)^*T*^). The leftmost matrix is *Cov*(*X*), the middle matrix is *Cov*(*X, Y*)*Cov*(*X,Y*)^*T*^, and the rightmost matrix is their element wise product *W*.

## 7 Spectral analysis

Assume a high dimensional set of variables *x_i_; i* = 1,…, *T*) where *T* >> *p*. We consider the case of a single functional module *M* with a subset of *p* variables from the high dimensional set. According to Theorem 6.1, we calculate *M* * = *diag*(*Cov*(*X *,Y*))*Var*(*X*\*)*diag*(*Cov*(*X*, Y*)) for the full set of *T* variables. Due to the arbitrary ordering of the variables, we will order the *p* module variables first for exposition and write *M*\* as a matrix with a positive upper diagonal block, *M*, and zeros elsewhere. By the Perron Theorem for positive matrices (Meyer (2000) ch. 8), the submatrix *M* contains the largest absolute eigenvalue *λ* > 0, with corresponding positive eigenvector *v*. Subsequently, *M*\* contains the largest absolute eigenvalue *λ* > 0 with sparse eigenvector (*v*, 0), where 0 denotes a vector of *T — p* zeros.

Our task is to extract a sparse eigenvector from matrix *W*, a noisy version of *M*\*, where the eigenvector is composed of either zeros or positive values. The nonzero eigenvector components are used to identify the module *M*. We introduce a novel method that follows naturally from the proceeding theoretical development. Note that an alternative to the more explanatory form (1) is *W* = *Z^T^Z*, where *Z* = *Hdiag*(*Cov*(*X*,Y*)) and *H* is the column-centered version of the *X*\* matrix. In summary, we calculate the transformed data matrix *Z*, recast the analysis as a sparse principal component analysis of *Z* using the algorithm from Sigg and Buhmann (2008), and then use the nonzero loadings from the sparse principal component analysis to select module elements.

The sparse principal component method from Sigg and Buhmann (2008) specifies sparsity by inputting the number of nonzero elements, *k*. We calculate the positive eigenvector for all feasible *k* (e.g. module sizes), and we select the sparsity setting *k* resulting in the optimally balanced solution. Balance may be quantified through the balance density 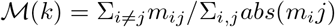 for a matrix in the form of (1). This can be motivated by an example where the matrix elements corresponding to the module of size *p* have values *r* > 0, and the non-module elements have values of 0. Thus, an effective sparse positive eigenvector algorithm will provide solutions

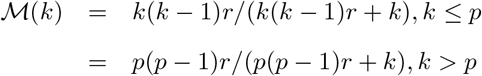

It then follows that 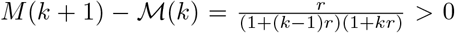 when *k* ≤ *p*, and 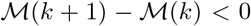 when *k* > *p*. This suggests that 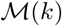 increases with respect to *k* for effective values of *k*. The decrease enlarges when negative elements are included. Thus, the maximum balance density may be used to tune the module size *k* in the Sigg-Buhmann algorithm for sparse principal component analysis.

## 8 Alternative characterization of balance

We provide a metric for evaluating the balance of a computed module based on an equivalent characterization of balance in an undirected signed graph: *a signed graph* G *is balanced if its vertices can be partitioned into subsets* A *and* B *where all edges within the subsets are positive and all edges connecting the vertices in different sets are negative* (Chartrand (1977), Ch. 8).

### Lemma 8.1

*With balanced A and B as defined above, all paths in the signed graph G connecting vertices in the same set are positive and all paths connecting a vertex in A to a vertex in B are negative.*

**Proof 8.1** *If a path in G connects vertices in A, then it must have zero or an even number of negative edges from A to B. Since all edges in A are positive, the path must be positive. Likewise for B. If a path in G connects a vertex in A to a vertex in B, then it must have zero or an odd number of negative edges from A to B. Since edges within A and B are positive, the path must be negative.*

For balanced semidirected graphs we can characterize correlations.

### Theorem 8.2

*For a balanced functional network G∪Y, G can be partitioned into two sets of variables A and B such that elements of A are positively correlated, elements of B are positively correlated, and correlations between A and B are negative. All elements of the same set have correlations with Y of the same sign, which is opposite for the two sets.*

**Proof 8.2** *By Lemma 8.1, the balanced graph G can be partitioned into sets A and B with positive paths connecting the vertices in the same set, and negative paths from one set to the other. By Theorem 5.2, variables in A are positively correlated, variables in B are positively correlated, and correlations between A and B are negative, proving the first statement. To prove the final statement, note that for a functional module G U Y there must be at least one variable j in G with an edge e: j → Y. Arbitrarily assume j ∈ A. By property 2 of Definition f.2, for each i ∈ A there is an i — j path. By Lemma 8.1, the i — j path is positive and so the i — j → Y path has the sign of e. By property 4 of Definition f.2, all i — Y paths for i ∈ A have the same sign as e, and by Theorem 5.3, Cov(i,Y) has the sign of e for all i ∈ A. Since G is connected, if k ∈ B then there exists a negative path to j by Lemma 8.1, and hence the k — j → Y path has an opposite sign from e. Thus, by Theorem 5.3, all variables in B have correlation with Y of sign opposite from e, proving the second statement of the theorem.*

Theorem 8.2 implies that for module elements *x_i_* and *x_j_*, the sign of *z_ij_* = *Cov*(*x_i_,y*) × *Cov*(*x_j_,y*) equals the sign of *Cov*(*x_i_, x_j_*). That is, if *x_i_* and *x_j_* reside in the same set, their correlation is positive and *sign*(*Cov*(*x_i_,y*)) = *sign*(*Cov*(*x_j_,y*)). If *x_i_* and *x_j_* reside in different sets, their correlation is negative and *sign*(*Cov*(*x_i_,y*)) ≠ *sign*(*Cov*(*x_j_,y*)). We flatten the upper diagonal elements of *Z* = (*z_i_,j*) and Σ into corresponding vectors of length *n*(*n* — 1)/2 and examine the agreement between the signs of the z-vector and the signs of the corelation vector. This can be viewed as the measurement of interrater reliablity of the two signs. Cohen’s Kappa measures agreement while considering the possibility of chance. A Cohen’s Kappa value less than or equal to 0 corresponds to no agreement/disagreement, and values greater than 0 correspond to agreement with perfect agreement equal to 1.0. We use Cohen’s Kappa to judge whether the sparse selection has captured agreeable variables according to Theorem 8.2, and hence whether they form a balanced functional module.

## 9 Simulation

### 9.1 Simulation model

A Single Input Module (SIM) consists of a set genes controlled by a single transcription factor (Milo et al. (2002)). Considerable experimental evidence shows that SIMs occur frequently (Lee et al. (2002), Milo et al. (2002)). For example, consider a SIM represented by the linear model for gene expression

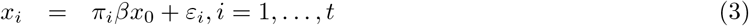

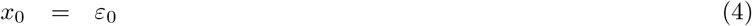

where *β* > 0, *π_i_* ∈ { — 1,1} and the *ε_i_, i* = 0,1,…, *t* are independent errors with mean 0 and variance 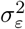. The covariance of all pairs of genes in this system are nonzero. The covariation among the *t* observed module genes are driven by a latent unobserved hub, *x*_0_. Letting *x_i_* and *x_j_* be two observed (non-hub) module genes, 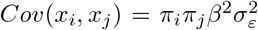 and 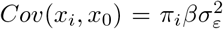. The non-hub variances equal 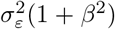, and the correlation between observed module genes *x_i_* and *x_j_* is 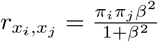.

We model the functional aspect of the pathway by letting the hub *x*_0_ determine an outcome variable *y* by the regression function

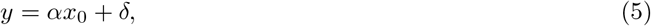

where *α* > 0 without loss of generality. Letting the variance of the error term *δ* in (5) be 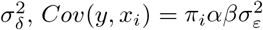 for a non-hub gene *x_i_*, 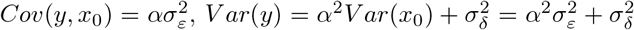, and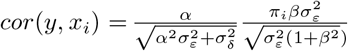.

A SIM is depicted in Figure 2. The unfilled circle *x*_0_ indicates that it is unobserved, while the blue vertices are observed variables. The hub *x*_0_ represents the latent “driver” of the pathway and the arrow from *x*_0_ to *y* represents the influence of the module on *y*. A negative sign *π_i_* on a link from *x*_0_ to *x_i_* indicates a positive change in *x*_0_ associates with an increase in *y* and suppression of *x_i_*, while a positive change indicates a promotion in *x_i_*.

**Figure 2:**
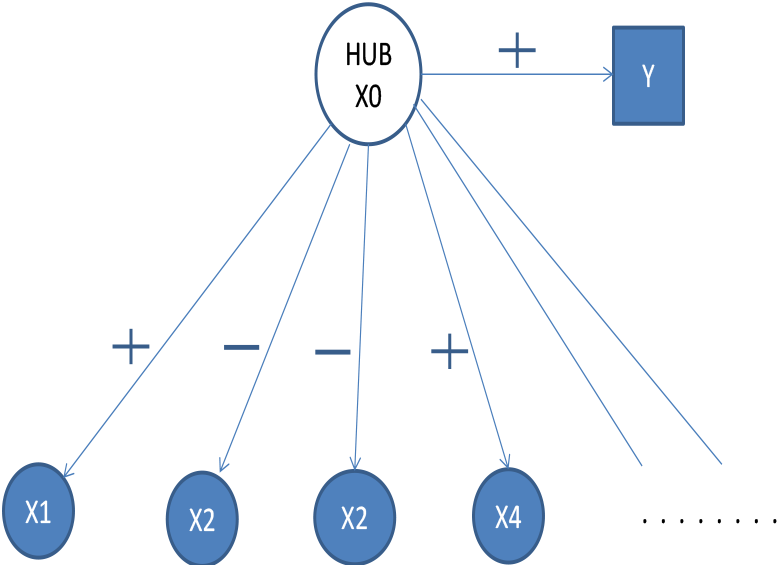
The graph of a SIM. The balance property is demonstrated by by tracing paths between vertices.

Tracing paths (Wright (1934)) in Figure 2 shows that a SIM is a balanced functional module. All predictor vertices positively correlated with Y are positively correlated with each other, and similarly for predictor vertices negatively correlated with Y. The path connecting a pair of vertices is negative if they have differing correlation with Y. Consequently, this simple latent variable model commonly used to illustrate biological phenomenon is actually a balanced module.

### 9.2 Experimental results

We simulated data containing 5 SIM modules with 20 vertices each and a set of 1000 irrelevant vertices, Δ. Only one of the modules is functional as per (5), and the other four are uncorrelated with the outcome variable *y*. We define signal-to-noise ratio (SNR) to be the ratio of *ασ_ε_* to *σ_δ_*, making SNR a ratio of the square root of the variance component in y attributed to the hub and the variance component in y attributed to the noise. We set model parameters to produce a specified SNR (SNR ∈ {.3,.5}) and intra-modular correlation (*r_m_* ∈ {.1, .2, .3, .4, .5, .6, .7}), for sample sizes of 100 and 400. In addition, we included 5 independent vertices with SNR equivalent to the hub node, for a total of 1,105 vertices.

We compare our method (bFMD) to three others. The first (W) was given by Miecznikowski et al. (2016) and operates on the matrix W defined previously. They set all negative elements of the matrix *W* to zero since the negative elements will correspond to unbalanced vertice pairs. The transformed matrix will then be matrix of elements corresponding to theoretical zeros and a highly positive block. Since the highly positive block will have a maximal sparse positive eigenvector according to the Perron Theorem, they use it to identify the module variables. The theoretical eigenvector has *p* positive elements and *N — p* zeros, so they partition the eigenvector elements into two clusters and take the cluster with a higher mean to be the module. Their method does not require specifying a sparsity parameter.

We also contrast our approach with Weighted Correlation Network Analysis (WGCNA), which is a two-step strategy of cluster analysis (modularity) followed by evaluating the identified clusters for the average correlation of the cluster members with the outcome variable (functionality) (Langfelder and Horvath (2008)). The functional information captured by the individual gene-outcome associations is not incorporated in the search for clusters at the first step. The basic approach can be implemented in a variety of ways by varying the similarity metric and the clustering method used, and it has been extended and highly developed (Zhang and Horvath (2005), Horvath (2011)) as part of a broad approach to network analysis. To implement it for our simulation study, we expressed similarity as absolute correlation and used PAM for the clustering method (Reynolds et al. (2006)). The clustering methods were set to obtain six clusters: one for the functional cluster, four for the nonfunctional clusters, and an additional cluster to capture other vertices. However, identifying the true number of clusters is difficult in practice. We computed the average significance for predicting *Y* for the genes in each cluster (the module significance), and the functional module was taken to be the most significant cluster.

In related work with a different objective, supervised Principal Components (SPC, Bair et al. (2006)) finds components composed of genes which are predictive of the outcome. In the first step, SPC selects a set genes highly associated with the outcome. Principal components are then computed for just those genes, and the resulting components are used for prediction. The number of selected genes is tuned using cross-validation to minimize the estimated out-of-sample prediction error using the principal component of the selected genes as the predictor. The components extracted by SPC are intended for prediction and not module identification. However, it is reasonable to consider using SPC for functional module discovery in the specific case that a single functional module influences the outcome since the genes selected would include the module genes and the principal component would load on the mutually correlated module genes. In this case, SPC exemplifies a two-step method which considers association with the outcome in the first step and co-expression in the second step, making it the reverse order compared to the WGCNA approach. For supervised principal components, we used the package “superpc” version 1.09 to calculate the first supervised principal component (Bair and Tibshirani (2004)).

We compared the balance based method bFMD with W, WGCNA and SPC. All methods are evaluated and compared in scenarios of coefficients with the same sign or random signs. In the simple case of coefficients with the same sign, we let *π_i_* = 1 in (3) for all module genes so that all intramodular correlations resulted in the same sign. For a more general network of coefficients with random signs, we randomly selected each *π* from the set { — 1,1} so that a module could consist of a mix of promoters and suppressors. 200 simulated samples were generated for each model under sample sizes of 100 and 400.

Given the underlying module used to simulate the data, we describe results for each method in terms of a) Sensitivity (the proportion of module elements included in the computed module), b) FDR (the proportion of the computed module that is falsely selected), c) the distribution of detected module size when the intramodule correlation is *r* = .4, and d) the Hamming distance between the detected module and the true module. Hamming distance is computed by defining *q* to be the binary vector which indicates selected variables by a procedure, and let *q*_0_ indicate the true module variables. The raw Hamming distance is the number of positions in disagreement across the two vectors. We report the distance normalized by the length of the vectors and express the proportion as a percentage, where 0 means perfect concordance and 100 means complete discordance. Hamming distance increases with both missed variables and false inclusions, making it a natural measure for the similarity between the computed module and the true module.

Figures 3 and 4 were generated for a sample size of 100 and *SNR* = 0.3 generated from randomly signed edges in the graph. Figure 3 shows the sensitivity and FDR of the methods. The method W from Miecznikowski et.al. selected the largest number of variables in the true module, followed by WGCNA and bFMD. The sensitivity of SPC plummets for *r* >. 2, presumably because the the goal of SPC is to select a parsimonious set of predictors, while the other methods employ colliniarity to define the module. bFMD by far results in the lowest FDR. The FDR curve for W is the most similar to bFMD, but it remains higher for all *r* values. The unacceptable FDR for WGCNA and SPC barely improves with higher intra-module correlation.

**Figure 3:**
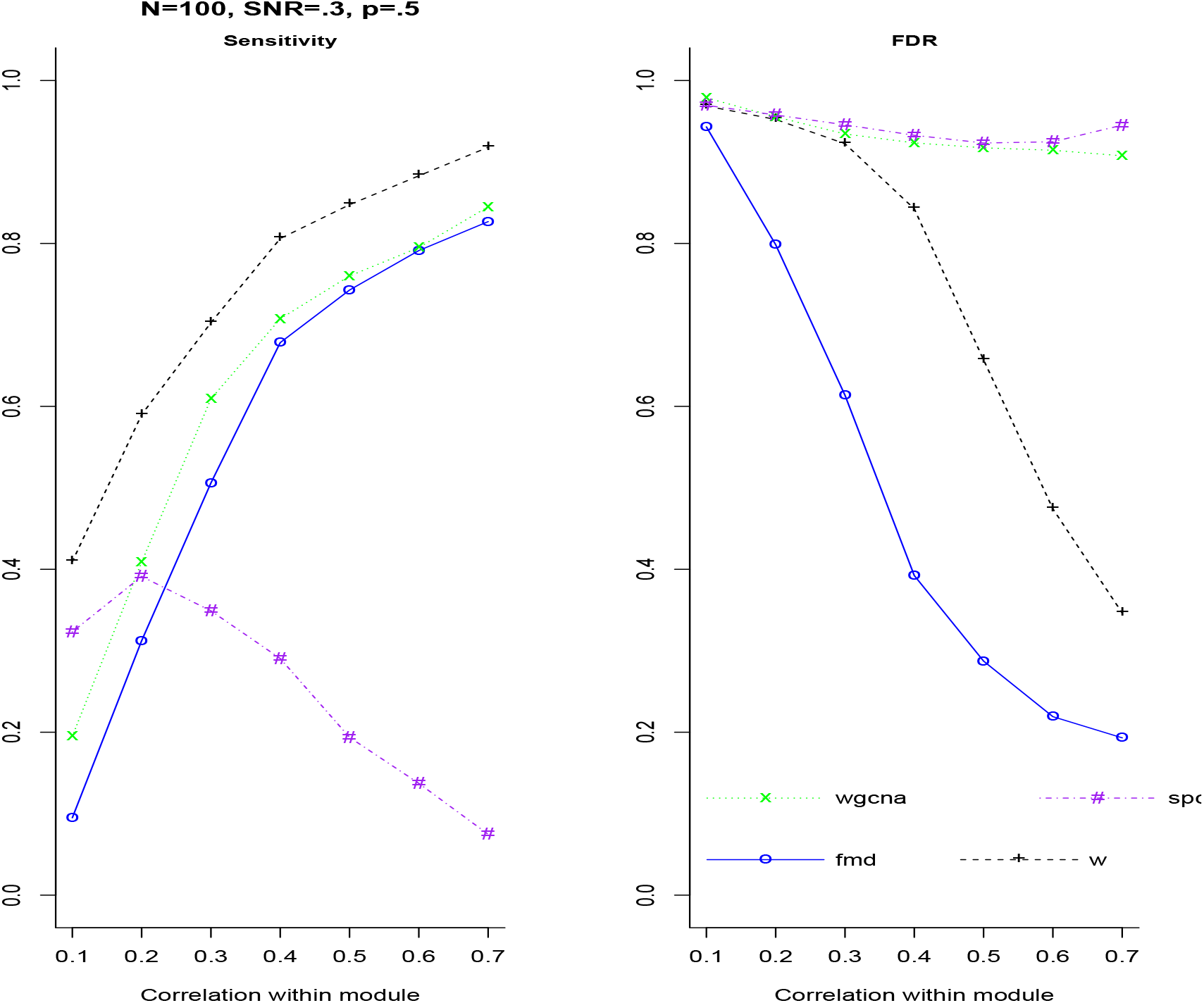
Sensitivity and FDR for *SNR* = 0.3 and sample size 100, generated with randomly-signed edges in the graph. bFMD is our balance-based module discovery method. W is the method from (Miecznikowski et al. (2016)), WGCNA is an implementation of Weighted Correlation Network Analysis (WGCNA) using k-means for 6 clusters (Langfelder and Horvath (2008)), and SPC is Sparse Principal Components (Bair et al. (2006)).

**Figure 4:**
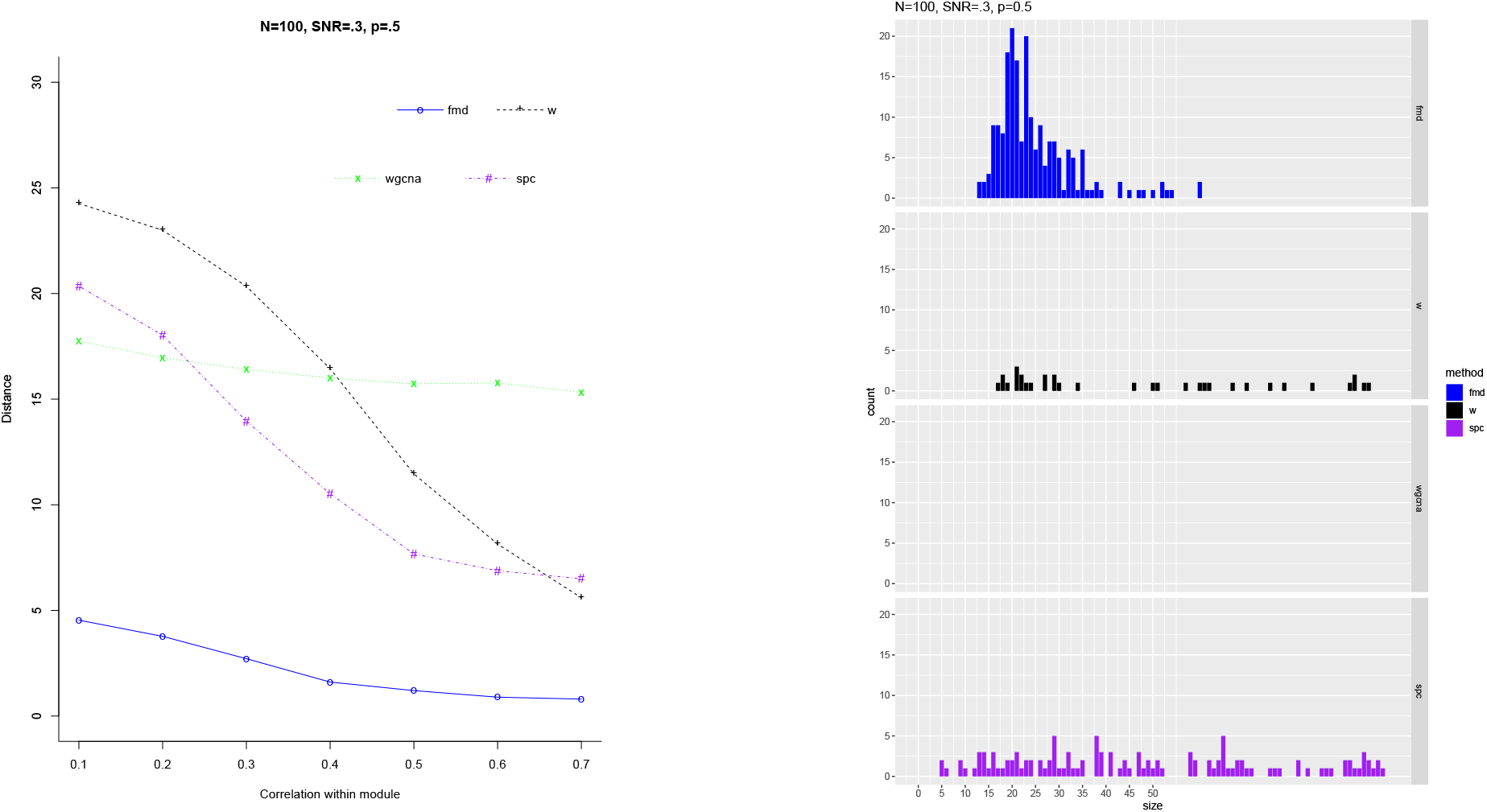
Hamming distance and distribution of module count for SNñ = .3 and sample size 100, generated with randomly signed edges in the graph. bFMD is our balance-based module discovery method. W is the method from (Miecznikowski et al. (2016)), WGCNA is an implementation of Weighted Correlation Network Analysis (WGCNA) using k-means for 6 clusters (Langfelder and Horvath (2008)), and SPC is Sparse Principal Components (Bair et al. (2006)).

Figure 4 compares the methods in terms of Hamming distance from the true module and shows the distributions of the module sizes found by the methods. bFMD has by far the lowest distance from the true module. The Hamming distance reflects both false positives and false negatives, and the superiority of bFMD suggests that the false discoveries heavily impact the distance measure for this data due to high dimensionality. WGCNA remains relatively unimproved by increasing intra-module correlation, suggesting that it is not tracking the true module. SPC performed better than W, which reflects the fact that parsimony tends to reduce the proportion of false negatives, ultimately influencing the Hamming distance in this high dimensional data.

The histgrams of module size are plotted for module size up to 100. It is also informative to examine the interquartile range of module size for each method: bFMD (19, 28), W (149, 258), WGCNA (178, 192), and SPC (37, 108). Note that since the minimum module size for WGCNA is 155, its histogram does not appear since the plot range only goes up to 100. bFMD has notably fewer extreme module sizes, and the selected modules from the other methods are often too large for useful interpretation. This explains the higher Hamming distance of SPC, WGCNA, and W.

When SNR is increased to .5, the performance of the methods improve due to the stronger signal, but the relative patterns remain similar, with the exception that the Hamming distance for W is greatly reduced. The interquartile ranges of module size become: bFMD (21, 25), W (21, 168), WGCNA (177, 191) and SPC (30, 317).

For increased sample size of 400 and *SNR* = .3, the characteristics of bFMD and W tend to converge. The sensitity of bFMD matches that of W for *r* ≥ .2, and the FDR and Hamming distance of W match that of bFMD for *r* ≥ .3. This is not suprising since they operate on a similar matrix, albeit in different ways. The Hamming distance for bFMD remains superior for all intra-module correlations.

To assess other module sizes, we ran similar simulations with the size of all modules increased to 50 with random signs and *SNR* = .3. Aside from the improved sensitivity in WGCNA, the patterns are consistent as in the simulations with module sizes of 20. Additionally, conclusions remain unchanged when repeating all of the simulations with entirely positive edges.

In summary, bFMD and W are superior to the other two methods. WGCNA suffers by not taking the association with Y into account when calculating clusters, which tend to be too big. SPC will select variables associated with *Y*, but those which are not correlated with other predictors will be missed. Only bFMD and W simultaneously consider both clustering and prediction criteria. W is more sensitive than bFMD although the difference diminishes greatly for larger intra-module correlations. This is due to W producing larger modules, thus decreasing the probability of missing an important variable. However, the higher sensitivity in W also increases its FDR compared to bFMD since W selects too many irrelevant variables. The results for W and bFMD tend to converge with increased sample size or SNR due to the higher signal strength. If investigators are very adverse to missing any important variable, they should use the W approach, but if they are interested in getting the most accurate representation and clearest interpretation of the module, bFMD should be the choice.

## 10 Example

The fourth leading cancer amongst women in the United States may be attributed to Uterine Corpus Endometrial Carcinoma (UCEC), according to (Levine et al. (2013)). In 2013, there were 50,000 new diagnosed UCEC cases and the cause of approximately 8000 deaths in the United States. UCEC was selected for assessment in the Pan-Cancer project within TCGA since achieving a sufficient sample size would be made easy due to its high incidence. Mutation, copy number, gene expression, DNA methylation, MicroRNA, and RPPA were obtained from tumor samples across 545 donors, and the data was made publicly available in the TCGA database (Weinstein et al. (2013)).

For this example, we will focus on discovering the subset of genes from the the gene expression dataset (X) that form a balanced module relating to percent tumor invasion (*Y*). The entire gene expression dataset within the TCGA-UCEC project contains 56457. To help reduce the amount of data in *X*, we will leverage prior research that shows mutations in MSH1, MSH2, MSH6, and PMS2 have been linked to Lynch Syndome and ultimately Endometrial Cancers Kempers et al. (2011). Since MSH2 and MSH6 both reside on chromosome 2 and have been shown to be associated with one another Schweizer et al. (2001), we will focus on chromosome 2 for the analysis. Percent tumor invasion was selected as the outcome for this example since it associates with tumor grade and patient survival (Boronow et al. (1984)).

Upon initial query of the TCGA-UCEC data, the clinical dataset (containing the information on percent tumor invasion) had 596 records and the gene expression dataset had 587 records. To clean the data, variable values within the gene expression dataset were averaged across any duplicate subject records, only the intersecting subject IDs across both the clinical and gene expression datasets were considered, and any subjects with missing information on percent tumor invasion were excluded. Subsequently, only the genes from chromosome 2 were considered and any genes with 0 variance were excluded. Furthermore, since the percent tumor invasion should range from 0 to 100, any patients with values outside of this range will be removed. Ultimately, the analytical set resulted in 469 subjects, each with a percent tumor invasion outcome and 3811 genes to consider for the functional module. In notational form, we now have a matrix *X* with 469 rows and 3811 variables (e.g. genes) and an outcome vector *Y* of length 469 (e.g. percent tumor invasion).

Analyses were done using bFMD, W, WGCNA and SPC. We describe the connectivity of estimated networks by the average correlation between module members, and the effect of the module on the outcome by the average correlation of module members with *Y*. For WGCNA we estimated that there are 6 clusters by inspecting the eigenvalues of the correlation matrix (Fallah et al. (2008)). Table 1 summarizes the results, and we note that bFMD out performs the other methods in every metric. While achieving the largest average module correlation and module effect, bFMD also returns the smallest module size, indicating a more tightly connected network. The value of Kappa for bFMD is .961, suggesting that the sparse eigenvector algorithm has produced a highly balanced module.

**Table 1:** Transcription data: Properties of estimated networks

|  | bFMD | W | WGCNA | SPC |
| --- | --- | --- | --- | --- |
| Module size | 187 | 680 | 795 | 248 |
| Module correlation | .23 | .19 | .05 | .12 |
| Module effect | .10 | .08 | .03 | .09 |

The 187 features selected by bFMD included the MSH2 and MSH6 genes which provides biological relevancy to the results. The W and WGCNA methods also selected the MSH2 and MSH6 genes, but they also selected many more genes which likely included noise due to the decreased module correlation and module effect. Despite the second smallest module size and moderate module correlation and module effects, SPC failed to select the MSH2 and MSH6 genes, weakening the biological interpretation of its selections. Figure 5 displays the selected module genes that are most negatively and positively correlated with MSH2 and MSH6.

**Figure 5:**
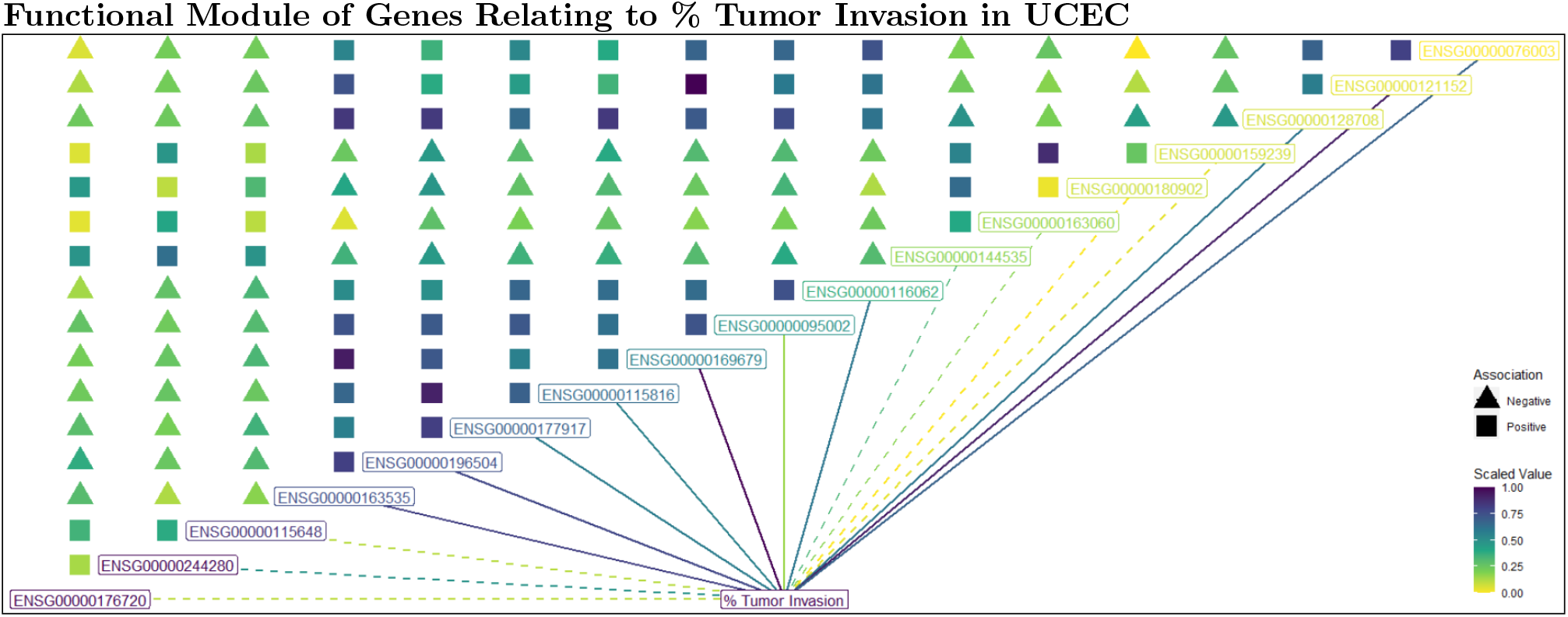
The selected genes obtained from bFMD on the gene expression from the TCGA-UCEC data that are most positively and negatively correlated with the biologically known genes MSH2 (ENSG00000095002) and MSH6 (ENSG00000116062). Since the full selected module is too large to display, only a subset of the selected genes is shown. The colored squares correspond to the absolute value of correlation between the genes within the module. In this example, all selected functional module genes are positively correlated with each other. The color of the labelled genes corresponds to the loading value obtained from the sparse principal component analysis, where the darker colors may be described as having a stronger relationship within the functional module. The dotted lines leading to the outcome label corresponds to the absolute value of correlation strength between the gene and the outcome, and the dotted lines (as opposed to solid) suggests that all genes within this functional module are negatively correlated with the outcome.

Table 2 shows the raw Hamming distances between the variable selection vectors of the four methods.

**Table 2:**
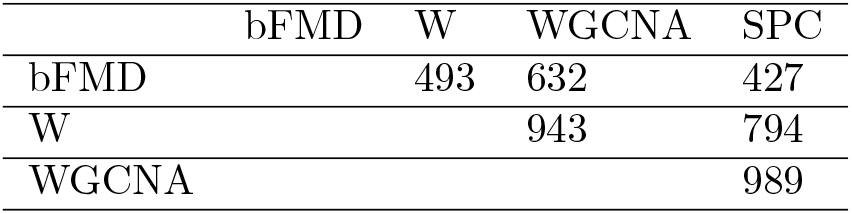
Transcription data: Hamming distance between network variable selection vectors of bFMD, W, WGCNA and SPC.

## 11 Summary and Discussion

The elements of an efficient biological mechanism may be expected to exhibit balanced effects which complement each other. The methods bFMD and W both operate on a matrix which highlights balanced sets of variables affecting an outcome variable as a positive submatrix. Of the two, bFMD most accurately identifies the set of module variables, as measured by the Hamming distance. Once the variables are extracted from a high-dimensional superset, detailed analysis of individual associations can be performed using existing methods for graphical model analysis (e.g. Wermuth and Cox (2013), Edwards (2000), Wermuth (2012), or Whittaker (2009)).

Our method is appropriate when linear relationships represent relevant aspects of the data. The theorems in this paper rely on the assumption of moderate conditional independence among variables and should be considered approximate for typical data sets. As with many statistical methods, strong multicollinearity potentially degrades performance.

In general, there could be multiple independent functional modules *M*_1_, *M*_2_,…, *M_K_*. In future work, deflation methods designed for sparse models such as (Mackey (2009)) will be investigated for cases with multiple processes operating.

## Acknowledgements

The author wishes to thank Dr. Xwei Chen for code used for the simulatons for this paper.

